# T4SE-XGB: interpretable sequence-based prediction of type IV secreted effectors using eXtreme gradient boosting algorithm

**DOI:** 10.1101/2020.06.18.158253

**Authors:** Tianhang Chen, Xiangeng Wang, Yanyi Chu, Dong-Qing Wei, Yi Xiong

## Abstract

Type IV secreted effectors (T4SEs) can be translocated into the cytosol of host cells via type IV secretion system (T4SS) and cause diseases. However, experimental approaches to identify T4SEs are time- and resource-consuming, and the existing computational tools based on machine learning techniques have some obvious limitations such as the lack of interpretability in the prediction models. In this study, we proposed a new model, T4SE-XGB, which uses the eXtreme gradient boosting (XGBoost) algorithm for accurate identification of type IV effectors based on optimal features based on protein sequences. After trying 20 different types of features, the best performance was achieved when all features were fed into XGBoost by the 5-fold cross validation in comparison with other machine learning methods. Then, the ReliefF algorithm was adopted to get the optimal feature set on our dataset, which further improved the model performance. T4SE-XGB exhibited highest predictive performance on the independent test set and outperformed other published prediction tools. Furthermore, the SHAP method was used to interpret the contribution of features to model predictions. The identification of key features can contribute to improved understanding of multifactorial contributors to host-pathogen interactions and bacterial pathogenesis. In addition to type IV effector prediction, we believe that the proposed framework can provide instructive guidance for similar studies to construct prediction methods on related biological problems. The data and source code of this study can be freely accessed at https://github.com/CT001002/T4SE-XGB.

## 1. Introduction

Different secretion systems have been found in bacteria that secret proteins into the extracellular environment. Gram-negative bacterial secretion can be categorized into eight types (from type I to type VIII), and the secreted proteins (also called effectors) play a vital role in bacterial pathogenesis and bacterium-host interactions. Some databases or web resource have been developed to store the experimentally validated effectors of Type III, IV, and VI secretion systems (An, et al., 2017; Bi, et al., 2013; Eichinger, et al., 2016; Li, et al., 2015). Type IV secretion system (T4SS) are protein complexes found in various species that deliver proteins into the cytoplasm of host cell and thus cause infection, such as whooping cough (Dorji, et al., 2018), gastritis, peptic ulcer and crown-gall tumor (Kuzmanovic, et al., 2018). Therefore, the identification of type IV secreted effector proteins (T4SEs) is a fundamental step toward understanding of the pathogenic mechanism of T4SS. There are a variety of experimental methods for identifying new T4SEs such as immunoblot analysis and pull-down assay (Cunha, et al., 2015). However, they are limited by both a *priori* knowledge about biological mechanisms and the sophisticated implementation of molecular experiments (Zeng and Zou, 2019). Furthermore, these experimental approaches are quite time-consuming and expensive. Instead, a large number of computational methods have been developed for prediction of T4SEs in the last decade, which successfully speed up the process in terms of time and efficiency. These computational approaches can be categorized into two main groups: the first group of approaches infer new effectors based on sequence similarity with currently known effectors (Chen, et al., 2010; Lockwood, et al., 2011; Marchesini, et al., 2011; Meyer, et al., 2013; Noroy, et al., 2019; Sankarasubramanian, et al., 2016) or phylogenetic profiling analysis (Zalguizuri, et al., 2019), and the second group of approaches involve learning the patterns of known secreted effectors that distinguish them from non-secreted proteins based on machine learning and deep learning techniques (Acici, et al., 2019; Ashari, et al., 2017; Burstein, et al., 2009; Chao, et al., 2019; Esna Ashari, et al., 2019; Esna Ashari, et al., 2019; Esna Ashari, et al., 2018; Guo, et al., 2018; Hong, et al., 2019; Li, et al., 2020; Lifshitz, et al., 2013; Wang, et al., 2019; Wang, et al., 2017; Wang, et al., 2014; Xiong, et al., 2018; Xue, et al., 2018; Yan, et al., 2020; Zou, et al., 2013). In the latter group of methods, Burstein et al. (Burstein, et al., 2009) worked on *Legionella pneumophila* to identify T4SEs and validated 40 novel effectors which were predicted by machine learning algorithms. Several features such as genomic organization, evolutionary based attributes, regulatory network attributes, and attributes specific to the *L. pneumophila* pathogenesis system were applied as input of the different machine learning algorithms: naïve Bayes, Bayesian networks, support vector machine (SVM), neural network and a voting classifier based on these four algorithms. Then, Zou et al. (Zou, et al., 2013) built the tool called T4EffPred based on the SVM algorithm with features such as amino acid composition (AAC), dipeptide composition (DPC), position specific scoring matrix composition (PSSM), auto covariance transformation of PSSM to identify T4SEs. Wang et al. (Wang, et al., 2014) constructed an effective inter-species T4SS effector prediction tool named T4SEpre, based on SVM by using C-terminal sequential and position-specific amino acid compositions, possible motifs and structural features. Later, Xiong et al. (Xiong, et al., 2018) used the same dataset as that of the previous study (Wang, et al., 2017) and developed a stacked ensemble classifier PredT4SE-Stack using various machine learning algorithms, such as SVM, gradient boosting machine, and extremely randomized trees. Wang et al. (Wang, et al., 2019) developed an ensemble classifier called Bastion4 which serves as an online T4SS effector predictor. They calculated 10 types of sequence-derived features. Then, the naïve Bayes, *k*-nearest neighbor, logistic regression, random forest, SVM and multilayer perceptron were trained and compared. Significantly improved predictive performance was achieved when they used the majority voting strategy based on the six classifiers where the PSSM-based features were used as input vectors. Recently, Ashari et al. developed the software called OPT4e (Esna Ashari, et al., 2019), which assembled various features used in prior studies to predict a set of candidate effectors for *A. phagocytophilum.* This tool yielded reasonable candidate effector predictions for most T4SS bacteria from the *Alphaproteobacteria* and *Gammaproteobacteria* classes.

Besides the traditional machine learning methods, deep learning is a new technology based on neural network architecture and has been successfully applied in wide range of applications in recent years (Deng, et al., 2020; Lv, et al., 2019; Ren, et al., 2019; Wu, et al., 2019; Yu, et al., 2018). Some researchers explored deep learning techniques to identify T4SEs based on protein sequences. Xue et al. (Xue, et al., 2018) proposed a deep learning method to identify T4SEs from protein sequences. The model called DeepT4 utilized a convolutional neural network (CNN) to extract T4SEs-related features from 50 N-terminal and 100 C-terminal residues of the proteins. This work provided the original idea about using the deep learning method. However, only few information of protein sequences can be extracted, which showed a slightly weaker performance compared with the Bastion4 (Wang, et al., 2019). Later, Açici et al. (Acici, et al., 2019) developed the CNN-based model based on the conversion from protein sequences to images using AAC, DPC and PSSM feature extraction methods. More recently, Hong et al. (Hong, et al., 2019) developed the new tool CNN-T4SE based on CNN, which integrated three encoding strategies: PSSM, protein secondary structure & solvent accessibility (PSSSA) and one-hot encoding scheme (Onehot), respectively. Compared with other machine learning methods, CNN-T4SE outperform all other state-of-the-art sequence-based T4SEs predictors. However, the less-than-optimal features analysis causes the limited deep learning for protein data and it is not straightforward to understand which features extracted from a given protein sequence drive the final prediction.

In this study, we proposed T4SE-XGB to predict type IV effectors using XGBoost based on sequence-derived features. To overcome the limitations of existing methods, we selectively summarized the features covered in previous studies and added some new features. The main strength of our method hinges on two aspects. On one hand, T4SE-XGB trained with features selected by the ReliefF algorithm significantly improved the overall performance on the benchmark dataset. On the other hand, T4SE-XGB uses a *post-hoc* interpretation technique: the SHAP method to demystify and explain specific features that led to deeper understanding of “black box” models.

## 2. Materials and Methods

The overall workflow of T4SE-XGB is shown in Figure 1, which is composed of five stages: Dataset Collection, Feature Extraction, Feature Selection, Model Construction and Model Interpretation. The detailed steps are described in the following sections.

**Figure 1.**
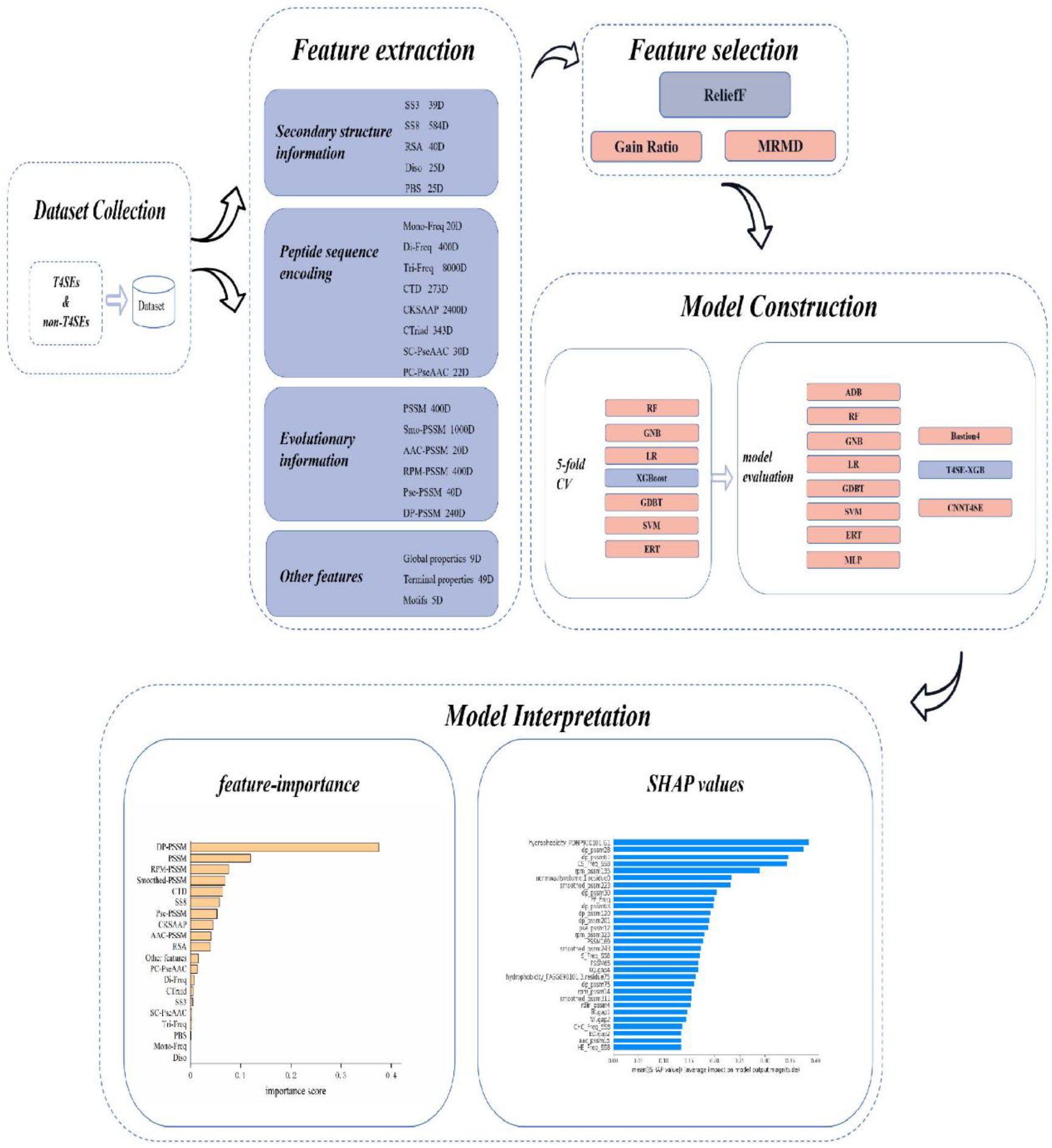
Overview workflow of T4SE-XGB. First, the benchmark dataset was collected. Next, 20 types of features were used to extract information from original protein sequences. Then, the ReliefF algorithm was employed to select optimal features. Five-fold cross validation test and independent test were set to verify the validation of the model. Finally, we not only used the vanilla XGBoost method to get feature importance, but also got SHAP values to realize the model interpretation.

### 2.1. Dataset

In our study, type IV secreted effectors and non-effectors were selected to build the benchmark dataset to construct the machine learning-based model for prediction of T4SEs. Our dataset was directly obtained from the recently published study (Wang, et al., 2019), which contained 420 T4SEs and 1262 non-T4SEs. The protein dataset was passed through a filter of >30% sequence identity to reduce sequence redundancy by the CD-HIT tool (Huang, et al., 2010). In the end, we got the final training dataset consisting of 365 T4SE and 1106 non-T4SEs, and the independent test dataset consisting of 20 T4SEs and 139 non-T4SEs.

### 2.2. Feature extraction

In this work, we took full advantage of features derived from protein sequences that former researchers have used and also added some novel features, which were used in other large-scale protein function prediction problems. We utilized the following four aspects of features to characterize protein sequences: secondary structure information (Zhu, et al., 2019), peptide sequence encoding (Yang, et al., 2019), evolutionary information and other features. Details about feature extraction are listed as below.

#### 2.2.1. Secondary structure information

(i) First, we used SCRATCH (Cheng, et al., 2005) to predict 3- and 8-state secondary structure (SS) information of amino acids of sequences and then *mono-* (1 state i.e. turn, strand or coil), *di-* (two consecutive states) and *tri*-state (three consecutive states) frequencies from a given protein sequence were extracted. (ii) The fraction of exposed residues (FER) with 20 different relative solvent accessibility (RSA) cutoffs (0% to 95% cutoffs at 5% intervals) and the FER by the average hydrophobicity of these exposed residues at different RSA cutoffs were calculated. (iii) DISOPRED (Ward, et al., 2004) can predict precise disordered region with annotated protein-binding activity. In the former study by Elbasir et al. (Elbasir, et al., 2019), they used DISOPRED to get 25 disordered features and 25 features of protein binding sites (PBS) in disordered regions.

#### 2.2.2. Peptide sequence encoding

(i) Frequencies of 20 amino acids, 400 di-peptides, 8000 tri-peptides were extracted from the protein sequences. (ii) The Composition, Transition and Distribution (CTD) feature represents the amino acid distribution patterns of a specific structural or physicochemical property in a protein or peptide sequence. Various types of physicochemical properties such as hydrophobicity, normalized Van der Waals Volume, polarity, polarizability, charge, secondary structures and solvent accessibility have been used for constituting the final feature vectors. (iii) The composition of *k-* spaced amino acid pairs (CKSAAP) feature calculates frequencies of amino acid pairs separated by any *k* (ranging from 0 to 5) residues. We use the default maximum value of *k* which is 5, and got a 2400-dimensional feature vector for one protein sequence. (iv) The Conjoint Triad descriptor (CTriad) considers the properties of one amino acid and its vicinal amino acids by regarding any three continuous amino acids as a single unit (Shen, et al., 2007). The occurrence that each triad appearing in the protein sequence is used to constitute a 343-dimensional vector after the amino acids are categorized into seven classes. (v) Pseudo amino acid composition analyses protein sequences about the physicochemical properties of the constituent amino acids. The final feature vectors include the global or long-range sequence order information. Series correlation pseudo amino acid composition (SC-PseAAC) (Chou, 2004) is a variant of PseAAC, which generates protein feature vectors by combining the amino acid composition and global sequence-order effects via series correlation. Parallel correlation pseudo amino acid composition (PC-PseAAC) (Chou, 2001), derived from PseACC, incorporates the contiguous local sequence-order information and the global sequence-order information into feature vectors of protein sequences.

The iFeature (Chen, et al., 2018) sever is capable of calculating and extracting different sequence-based, structural and physiochemical features derived from protein sequences. The BioSeq-Analysis2.0 (Liu, et al., 2019) sever was employed to generate modes of pseudo amino acid compositions (such as SC-PseAAC and PC-PseAAC) for protein sequences.

#### 2.2.3. Evolutionary information

(i) PSSM of a protein sequence can be obtained in the form of L*20 matrix (L is the amino acid length). PSSM represents the evolutionary, residue and sequence information features of input proteins. In our study, we got 400 feature vectors from the original PSSM profile by summing rows corresponding to the same amino acid residue. (ii) Smoothed-PSSM (Cheng, et al., 2008) transformed from the standard PSSM encodes the correlation or dependency from surrounding residues, which significantly enhanced the performance of RNA-binding site prediction in proteins.

The Smoothed-PSSM profile considered the first 50 amino acids starting from the protein’s N-terminus to form a vector with the dimension 1000. (iii) AAC-PSSM (Liu, et al., 2010) represents the correlation of evolutionary conservation of the 20 residues between two positions separated by a predefined distance along the sequence and successfully converts a protein into a fixed length feature vector with dimension 20. It reveals the possibility of the amino acid residues mutated to different types during the evolution process. (iv) RPM-PSSM (Jeong, et al., 2011) filters all entities with values of less than 0 from the PSSM matrix by using the residue probing method, in which each amino acid is regarded as a probe corresponds to a particular column in the PSSM profiles, and the negative values were set to 0. For this method, original PSSM matrix finally transformed into the 20*20 matrix and can be constructed into a 400-dimensional vector. (v) Pse-PSSM (Chou and Shen, 2007) is similar to PseAAC and encodes the PSSM of proteins with different lengths using a uniform length matrix. (iv) DP-PSSM (Juan, et al., 2009), a protein descriptor based on similarity, gets the hidden sequential order information and can avoid cancellation of positive or negative terms in the average process. As a result, we obtained a 400-dimensional vector for each sequence.

These PSSM-based features were achieved using the bioinformatics tool called POSSUM (Wang, et al., 2017), including the original PSSM profiles, smoothed-PSSM, AAC-PSSM, RPM-PSSM, Pse-PSSM and DP-PSSM. All PSSM-based features used default parameters the website provided: smoothing window=7 and sliding window=50 for smoothed-PSSM, ξ=1 for Pse-PSSM, and α=5 for DP-PSSM.

#### 2.2.4. Other features

(i) Global properties of the protein were calculated such as sequence length, molecular weight, and total hydropathy et al. (ii) Terminal properties like the frequencies of 20 amino acid types in the 50 amino acids at the C-terminus or N-terminus used in previous studies were also calculated (An, et al., 2018; Wang, et al., 2019; Wang, et al., 2017; Zeng and Zou, 2019). The frequencies of di-peptides at the C-terminus, like SS, KE, EE, EK, AA, AG and LL involved in former studies have shown variances between effectors and the non-effectors were also calculated (Zou and Chen, 2016; Zou, et al., 2013). (iii) We also searched for several types of protein motifs including nuclear localization signals (NLS), E-Block (EEXXE motif), conserved EPIYA motifs (EPIYA_CON), hypothetical EPIYA motifs (EPIYA_HYS) and Prenylation Domain (CaaX motif) that have been proposed and extracted before (Esna Ashari, et al., 2019; Esna Ashari, et al., 2018; Noroy, et al., 2019).

### 2.3. Feature normalization

Normalization is a scaling technique in which values are shifted and rescaled so that they fall into the same numeric interval. Having features on a similar scale helps the gradient descent converge more quickly towards the minima. The following formula can be used to normalize all feature values and end up ranging between 0 and 1, which is known as Min-Max scaling:

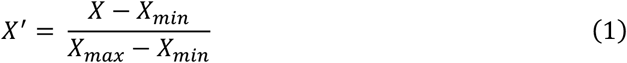

Here, *X_max_* and *X_min_* are the maximum and the minimum values of the feature respectively. We imported the MinMaxScalar package from the python scikit learn library to calculate the normalized values.

### 2.4. Feature selection

The purpose of dimensionality reduction or feature selection is to reduce the computational time and complexity of the prediction model, and also to provide more insights into the data abundance (Basith, et al., 2020; Govindaraj, et al., 2020; He, et al., 2018; Jing, et al., 2019; Kang, et al., 2019; Li, et al., 2020; Liu, et al., 2019; Manavalan, et al., 2018; Shi, et al., 2019; Su, et al., 2020; Tang, et al., 2018; Xiong, et al., 2012; Xiong, et al., 2019; Zhang, et al., 2020). It is indispensable to reduce dimensionality to remove redundant features so that we can reserve the important ones.

#### 2.4.1. Gain ratio

The gain ratio algorithm based on information theory can be used to deal with oversized feature sets (Shannon, 1948). We used the gain.ratio function from the R package named FSelector. The algorithm finds weights of discrete attributes based on their correlation with continuous class attribute.

#### 2.4.2. ReliefF

The ReliefF algorithm is an improvement of Kononenko’s standard Relief algorithm (Kira and Rendell, 1992). In this work, the ReliefF algorithm was implemented by the ReliefFexpRank function in the attrEval method from R package named CORElearn (Yu, et al., 2019). Rank of nearest instance is determined by the increasing (Manhattan) distance from the selected instance and the *k* nearest instances have weight exponentially decreasing with increasing rank. This is a default choice for methods taking conditional dependencies among the attributes into account.

The ReliefF algorithm fully considers the correlation between features and labels, in order to effectively remove unnecessary features after updating the feature weights according to the degree of correlation. The higher the weight value, the stronger the classification ability of the feature. The weight W of each feature is defined as:

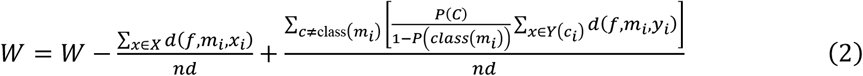

where *d* means samples with the nearest distance *m_i_* from each category selected by the ReliefF algorithm first, *f* means a certain feature, *n* means the number of samples, and *d*(*f,m_i_,x*) means the distance between sample *X* and sample *Y* for a certain feature *f*.

#### 2.4.3. Maximum-Relevance-Maximum-Distance

The Maximum-Relevance-Maximum-Distance (MRMD) algorithm uses the Pearson’s correlation coefficient to measure the relevance between features in a subset. The Pearson correlation coefficient shows the degree of relationship between features and labels. Besides, Euclidean distance, cosine similarity and Tanimoto coefficient are utilized to calculate the redundancy between features in a subset. In the end, the MRMD algorithm selects features which have strong correlation with labels and have lowest redundancy between features (Zou, et al., 2016).

### 2.5. Extreme gradient boosting

Extreme gradient boosting also named XGBoost (Chen and Guestrin, 2016) is an optimized distributed gradient boosting algorithm designed to be highly efficient, flexible and portable (Wang, et al., 2019). XGBoost based on decision tree ensembles consists of a set of classification or regression trees. It uses the training data (with multiple features) *x_i_* to predict a target variable *y_i_*. To begin with, the objective function is defined as:

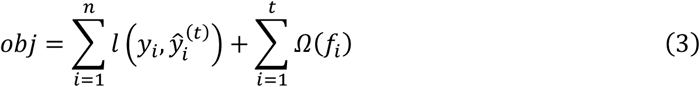

where *n* is the number of trees, *l* is the training loss function, Ω is the regularization term.

Then, the XGBoost takes the Taylor expansion of the loss function up to the second order and removes all the constants, so the specific objective at step t becomes:

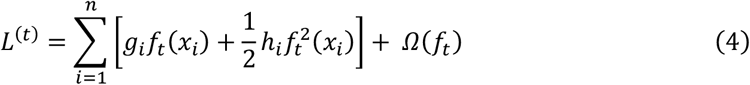

where the *g_i_* and *h_i_* are defined as

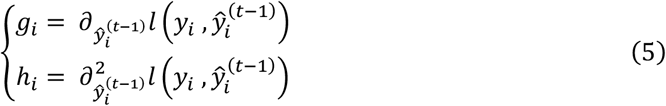

The value of the objective function only depends on *g_i_* and *h_i_*. It can optimize every loss function, including logistic regression and pairwise ranking.

The traditional treatment of tree learning only emphasized the improved impurity, while the complexity control was left to heuristics. Chen et al. (Chen and Guestrin, 2016) formally defined the complexity of the tree *Ω*(*f*) to obtain regularization, and the loss function in the *t*-th tree finally can be rewritten as:

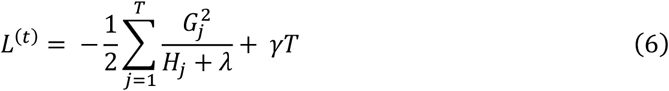

where the *G_j_* and *H_j_* are defined as

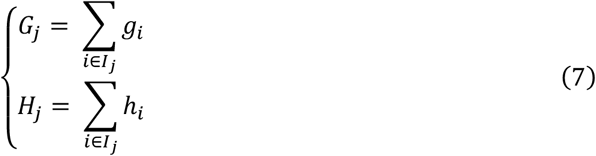

*I_j_* is the sample set divided into the *j*-th leaf node according to the decision rules for a given tree. The formula (6) can be used as the score value to evaluate the quality of a tree. They also defined the score it gains when a leaf is split into two leaves:

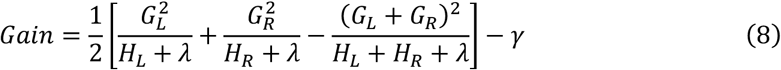

This formula is composed of the score on the new left leaf, the score on the new right leaf, and the score on the original leaf and regularization on the additional leaf. We can find the best split efficiently by the maximum value of *Gain* through a scan from left to right to get all possible split solutions.

XGBoost with many optimization techniques is able to solve problems using far fewer resources. It is simple to parallel and can greatly enhance the program efficiency with a fast model exploration. More details about XGBoost are given in (Chen and Guestrin, 2016).

## 2.6. Performance evaluation

In this work, confusion matrix obtained after prediction contains four units: true positive (TP), false positive (FP), false negative (FN) and true negative (TN). In order to evaluate the overall predictive performance of different classification models, we used metrics such as Sensitivity (SE), Specificity (SP), Precision (PRE), Accuracy (ACC), F-score and Matthew’s correlation coefficient (MCC) to evaluate the model. They have been widely used in previous studies (Al-Ajlan and El Allali, 2019; Cheng, et al., 2020; Chu, et al., 2019; Hasan, et al., 2020; Jing and Dong, 2017; Lin, et al., 2019; Liu, et al., 2020; Manavalan, et al., 2019; Yue, et al., 2020; Zhang, et al., 2019; Zhang, et al., 2019; Zhang, et al., 2020; Zhu, et al., 2019), with a higher value indicating better performances. The performance metrics can be defined as follows:

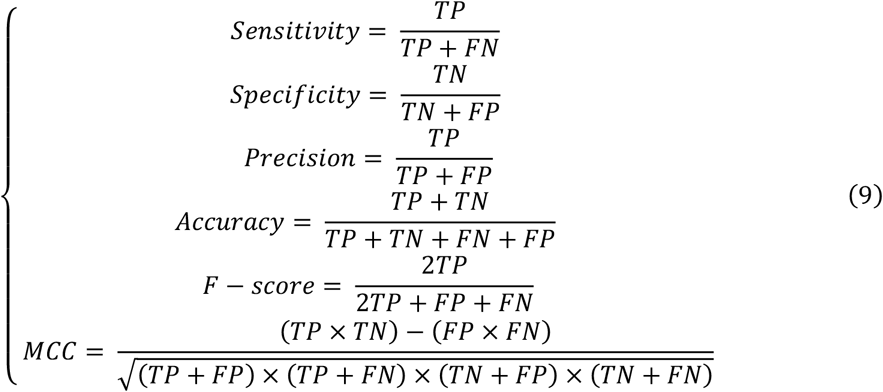

## 3. Results and Discussion

### 3.1. Comparison of different features and their combinations on the training data set

In this section, we evaluated the predictive power of the individual types of features and their combinations using the XGBoost classification algorithm by the 5-fold cross validation (CV) method on the training data set. In 5-fold CV, the training data set was randomly divided into five subsets. XGBoost were trained by four subsets and the remaining one was used to evaluate the performance of the model. All steps were repeated five times. The average of the performance measures such as ACC and SE of the training set were calculated and the results are shown in Table 1. It can be seen that some individual feature types based on PSSM have higher overall prediction power on the training data set. This observation indicates that the features based on PSSM have better performance in the prediction of T4SE when compared with other types of features.

**Table 1.** Comparison of predictive power of different features on the training data set by 5-fold cross validation test

| Feature Types | ACC (%) | SE (%) | PRE (%) | F-score | MCC |
| --- | --- | --- | --- | --- | --- |
| ss3 | 79.94 | 43.84 | 64.52 | 0.5209 | 0.4125 |
| ss8 | 81.44 | 48.77 | 68.57 | 0.5640 | 0.4650 |
| RSA | 84.23 | 58.90 | 72.80 | 0.6497 | 0.5556 |
| Diso | 79.81 | 35.62 | 68.16 | 0.4650 | 0.3856 |
| PBS | 79.47 | 36.16 | 65.74 | 0.4643 | 0.3760 |
| Mono-Freq | 85.18 | 60.00 | 75.54 | 0.6669 | 0.5809 |
| Di-Freq | 84.36 | 53.97 | 76.16 | 0.6302 | 0.5486 |
| Tri-Freq | 80.01 | 30.41 | 73.85 | 0.4301 | 0.3822 |
| PSSM | 92.11 | 78.08 | 88.99 | 0.8305 | 0.7833 |
| smoothed-PSSM | 88.51 | 70.14 | 81.03 | 0.7512 | 0.6806 |
| AAC-PSSM | 90.69 | 74.52 | 86.27 | 0.7976 | 0.7425 |
| RPM-PSSM | 91.98 | 77.26 | 89.32 | 0.8262 | 0.7797 |
| Pse-PSSM | 92.59 | 81.37 | 88.13 | 0.8446 | 0.7984 |
| DP-PSSM | 92.79 | 81.37 | 88.95 | 0.8486 | 0.8039 |
| CKSAAP | 84.02 | 51.78 | 76.31 | 0.6153 | 0.5359 |
| CTD | 87.70 | 67.12 | 80.24 | 0.7296 | 0.6562 |
| CTraid | 82.53 | 49.86 | 71.04 | 0.5816 | 0.4909 |
| SC-PseAAC | 85.66 | 61.92 | 75.96 | 0.6811 | 0.5958 |
| PC-PseAAC | 85.52 | 61.10 | 76.05 | 0.6769 | 0.5913 |
| Other features | 84.09 | 54.52 | 74.62 | 0.6288 | 0.5422 |
| All features | 93.95 | 81.92 | 93.04 | 0.8698 | 0.8346 |

The combination of different features could depict protein sequences in a more comprehensive manner (Wang, et al., 2019). As illustrated in Table 1, using the combined features yield the ACC of 93.95% and the MCC of 0.8346, which are both higher than other PSSM-based features. In summary, compared with single feature-based models, the combination of all features achieved consistently better performance.

### 3.2. Comparison of three feature selection methods on the training data set

In this section, three kinds of feature selection methods were compared on the training data set by the 5-fold cross validation test. They are gain ratio algorithm (Shannon, 1948), maximum relevance–maximum distance (MRMD) (Zou, et al., 2016) and ReliefF algorithm (Kira and Rendell, 1992),. The ACC of different number of dimensions were obtained and compared by using different feature selection algorithms to select most useful features. As shown in Table 2, when the MRMD algorithm was used for dimensionality reduction on the training data set, the highest ACC value was 92.93%. The gain ratio algorithm achieved ACC of 93.81% on the training data set. By comparing the prediction accuracy of three methods in different dimensions, it can be found that the ReliefF algorithm achieved the highest ACC value, 94.42% when the dimension was 1000, obviously better than the models using all original features.

**Table 2.** Comparison of different feature selection methods with different dimensions of the selected features on the training data set by 5-fold cross validation test

| Method | 500 | 600 | 700 | 800 | 900 | 1000 | 1100 | 1200 | 1300 | 1400 | 1500 |
| --- | --- | --- | --- | --- | --- | --- | --- | --- | --- | --- | --- |
| GainRatio | 92.79 | 92.73 | 93.34 | 93.34 | 93.41 | 93.47 | 93.34 | 93.54 | 93.60 | 93.41 | 93.81 |
| MRMD | 92.18 | 92.45 | 92.32 | 92.52 | 92.18 | 92.66 | 92.39 | 92.93 | 92.86 | 92.25 | 92.93 |
| ReliefF | 93.74 | 93.95 | 93.61 | 93.74 | 94.36 | 94.42 | 94.22 | 94.09 | 93.95 | 94.02 | 94.08 |

Therefore, the ReliefF algorithm can effectively eliminate redundant variables and improve prediction accuracy. In the following sections, the ReliefF algorithm was used for dimensionality reduction.

### 3.3. Comparison of different classification algorithms on the training data set

In order to objectively validate the prediction power of the XGBoost algorithm, we compared the performance of this algorithm with other classification algorithms by using the 5-fold cross validation on the training data set. Based on the optimal set of features, other classification algorithms such as Random Forests (RF) (Zhang, et al., 2020), naïve Bayes (NB), Logistic Regression (LR), Gradient Boost (GDBT), support vector machine, *k*-nearest neighbor (KNN), Extremely randomized trees (ERT) and Multi-layer Perceptron (MLP) were all trained and compared. The grid search method was employed in this work to optimize hyper-parameters for each classifier (Shan, et al., 2019). For each ML classifier, we obtained the best hyper-parameter combination based on the highest accuracy by the 5-fold cross validation. Table 3 shows the comparison results of XGBoost with other classification algorithms on the training data set by 5-fold cross validation.

**Table 3.** Comparison of different classification algorithms on the training data set by 5-fold cross validation test

|  | ACC (%) | SE (%) | PRE (%) | F-score | MCC |
| --- | --- | --- | --- | --- | --- |
| NB | 90.89 | 84.11 | 80.76 | 0.8207 | 0.7631 |
| ML | 92.32 | 82.47 | 86.10 | 0.8409 | 0.7920 |
| LR | 93.00 | 83.01 | 88.28 | 0.8539 | 0.8101 |
| KNN | 93.20 | 80.82 | 91.20 | 0.8544 | 0.8148 |
| RF | 93.27 | 80.27 | 91.94 | 0.8554 | 0.8163 |
| ERT | 93.54 | 80.55 | 92.76 | 0.8604 | 0.8235 |
| GDBT | 93.81 | 84.11 | 90.55 | 0.8710 | 0.8323 |
| SVM | 94.36 | 83.56 | 93.28 | 0.8794 | 0.8466 |
| XGB | 94.42 | 83.01 | 94.02 | 0.8803 | 0.8481 |

As shown in Table 3, the ACCs of different classifiers were falling within the range from 90.89% to 94.42%, and their MCCs were ranging from 0.76 to 0.84 on the training data set. The results showed that XGBoost achieved the best performance, where the ACC, F-score and MCC were significantly higher than the other classifiers. All in all, the XGBoost algorithm performs better than the other machine learning-based methods when applied on the training data set.

### 3.4. Comparison with other classification algorithms and existing methods on the independent test data set

To further validate the performance of the proposed model in the real test, we compared the performance of our T4SE-XGB model with other classification algorithms and several state-of-the-art methods on the independent data set. The performance results of these methods are provided in Table 4. To make a fair comparison, the same independent data set, which consists of 20 T4SEs and 139 non-T4SEs, was used for all models.

**Table 4.** Comparison of different classification algorithms and the state-of-the-art methods on the independent data set

| Model | TP | FN | TN | FP | ACC (%) | SE (%) | SP (%) | PRE (%) | F-score | MCC |
| --- | --- | --- | --- | --- | --- | --- | --- | --- | --- | --- |
| SVM | 19 | 1 | 134 | 5 | 96.23 | 95.00 | 96.40 | 79.17 | 0.8636 | 0.8467 |
| LR | 19 | 1 | 131 | 8 | 94.34 | 95.00 | 94.24 | 70.37 | 0.8085 | 0.7882 |
| NB | 19 | 1 | 126 | 13 | 91.19 | 95.00 | 90.65 | 59.38 | 0.7308 | 0.7084 |
| GDBT | 19 | 1 | 131 | 8 | 94.34 | 95.00 | 94.24 | 70.37 | 0.8085 | 0.7882 |
| RF | 19 | 1 | 132 | 7 | 94.97 | 94.96 | 95.68 | 73.08 | 0.8261 | 0.8066 |
| ERT | 19 | 1 | 134 | 5 | 96.23 | 95.00 | 96.40 | 79.17 | 0.8636 | 0.8467 |
| KNN | 20 | 0 | 128 | 11 | 93.08 | 100.0 | 92.09 | 64.52 | 0.7843 | 0.7708 |
| ML | 18 | 2 | 129 | 10 | 92.45 | 90.0 | 92.81 | 64.29 | 0.7500 | 0.7209 |
| Bastion4 | 20 | 0 | 133 | 6 | 96.23 | 100.0 | 95.68 | 76.92 | 0.8696 | 0.8579 |
| CNNT4SE(PSSSA) | 14 | 6 | 138 | 1 | 95.60 | 70.00 | 99.28 | 93.33 | 0.8000 | 0.7860 |
| CNNT4SE(Onehot) | 14 | 6 | 139 | 0 | 96.23 | 70.00 | 100.0 | 100.0 | 0.8235 | 0.8192 |
| CNNT4SE(PSSM) | 20 | 0 | 138 | 1 | 99.37 | 100.0 | 99.28 | 95.24 | 0.9756 | 0.9724 |
| CNNT4SE(VOTE 2/3) | 16 | 4 | 139 | 0 | 97.48 | 80.00 | 100.0 | 100.0 | 0.8889 | 0.8818 |
| T4SE-XGB | 19 | 1 | 136 | 3 | 97.48 | 95.00 | 97.84 | 86.36 | 0.9048 | 0.8916 |

Among these machine learning-based methods, the results showed that our T4SE-XGB model achieved the overall best performance with an ACC of 97.48%, F-value of 90.48% and MCC of 0.8916, followed by the state-of-the-art machine learning model called Bastion4 (Wang, et al., 2019), which achieved 96.23% on ACC, 86.96% on F-value and 0.8579 on MCC. Moreover, the T4SE-XGB trained by fewer training samples also gets more stable prediction performance than the deep learning-based method named CNN-T4SE (VOTE 2/3), which takes the majority votes of the three best-performing convolutional neural network-based models (CNN-PSSM, CNNPSSSA and CNN-Onehot). The CNN-PSSM, a deep learning-based model based on PSSM features, achieved the best results. However, it gets two less false positive and one less false negative when compared with our model.

In summary, there is a consistent observation (from the results obtained from the 5-fold cross validation test and independent test) that our T4SE-XGB model achieved higher performance in terms of sensitivity, specificity, accuracy and MCC on both the training data set and independent data set.

### 3.5. Model interpretation

#### 3.5.1. Estimation of feature importance by XGBoost

As a tree-based non-linear machine learning technique, XGBoost can exploit the interactions between the engineered features. In contrast to black-box modeling techniques such as SVM, ANN, CNN, the XGBoost algorithm can easily obtain feature importance scores for all input features. XGboost can also obtain the importance score efficiently based on the frequency of a feature which is used to split data or according to the average gain a feature brings when it was used during node splitting across all established trees. For the optimal set of features constructed on the benchmark dataset, the importance of each feature during training is the sum of information gained when used for splits (tree branching).

The total feature importance contribution of all features according to their feature types are shown in Table 5 and Figure 2. We can see that the DP-PSSM feature gets the maximum value of the importance score which is 0.3758. This may mean that the DP-PSSM feature is more important. Besides, the PSSM feature which incorporated evolutionary information has the importance score of 0.1199, followed by other features based on the transformation of the standard PSSM profile, such as RPM-PSSM and Smoothed-PSSM. There are also other features showing high importance. For example, CTD accounts for 6.46% of all feature importance score. SS8 makes up 5.84% of the total variable importance.

**Figure 2.**
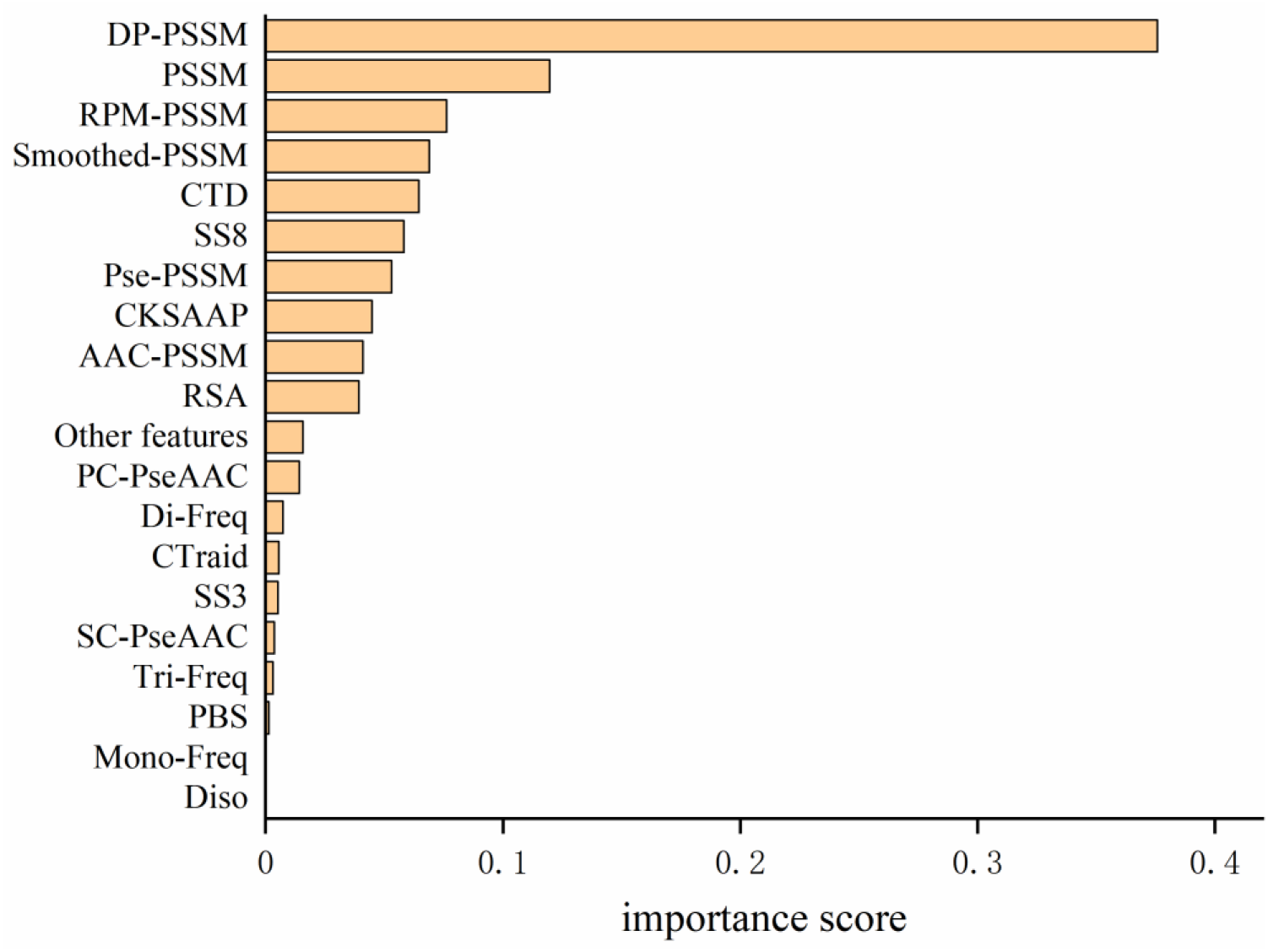
Comparison of importance percentages grouped by feature types for the T4SE-XGB model.

**Table 5.** Importance percentages grouped by feature types for the T4SE-XGB model

| Feature name | Importance score |
| --- | --- |
| DP-PSSM | 0.3758 |
| PSSM | 0.1199 |
| RPM-PSSM | 0.0764 |
| Smoothed-PSSM | 0.0690 |
| CTD | 0.0646 |
| SS8 | 0.0584 |
| Pse-PSSM | 0.0532 |
| CKSAAP | 0.0449 |
| AAC-PSSM | 0.0411 |
| RSA | 0.0394 |
| Other features | 0.0158 |
| PC-PseAAC | 0.0143 |
| Di-Freq | 0.0075 |
| CTraid | 0.0057 |
| SS3 | 0.0053 |
| SC-PseAAC | 0.0038 |
| Tri-Freq | 0.0032 |
| PBS | 0.0015 |
| Diso | 0 |
| Mono-Freq | 0 |

#### 3.5.2. Model interpretation by SHAP

SHAP (SHapley Additive exPlanations), a unified framework for interpreting predictions, assigns each feature an importance value for a particular prediction (Lundberg and Lee, 2017) and improves the interpretability of tree-based models such as random forests, decision trees, and gradient boosted trees (Lundberg, et al., 2020; Lundberg, et al., 2018). SHAP is based on the game theoretically optimal Shapley values that can be calculated as below (Lipovetsky and Conklin, 2001):

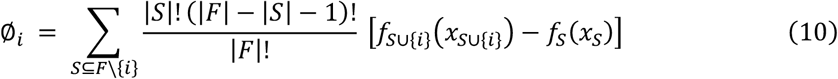

where *F* is the set of all features, *S* is a subset of the features used in the model without the *i^th^* feature, *x* is the feature vector of the instance to be explained. SHAP comes with many global interpretation methods based on aggregations of Shapley values. More detailed description of the SHAP method is available in (Lundberg and Lee, 2017).

The SHAP method has the ability to provide interpretable predictions and also overcomes limitation that the feature importance scores obtained from XGBoost model, which is in lack of directivity, and is unable to correspond to specific eigenvalues. Figure 3(A) is the standard bar-chart based on the average magnitude of the Shapley values over all training samples. The higher value indicates higher feature importance. It can be seen that DP-PSSM has the largest number of important features, accounting for 7, among the 30 most important features. Meanwhile, other features based on PSSM also form the majority. Among them, the hydrophobicity_PONP930101.G1 coming from the feature unit of CTD can be obviously identified as the most important. Hydrophobicity_PONP930101 is one physicochemical attribute based on the main clusters of the amino acid indices of Tomii and Kanehisa (Tomii and Kanehisa, 1996). The hydrophobicity_PONP93-0101.G1={ N(r)/N, r ∈ {KPDESNQT}} represents the global compositions (percentage) of polar residues of the protein under the hydrophobicity_PONP930101 attribute (Chen, et al., 2018). Several studies have suggested that type IV effector proteins exhibited some specificities in regard to amino acid frequency. Zou et al. (Zou, et al., 2013) calculated the ACC and the variance in their dataset. They found that Asn (N), Glu (E) and Lys (K) have higher compositions in type IVB effectors than non-effectors, and Ala (A), Glu (E) and Ser (S) have higher compositions in type IVA effectors than non-effectors. Some polar amino acids, such as Asp (D), Cys (C) and His (H), have small differences between secreted proteins and non-secreted proteins. Similarly, The Mann–Whitney U-test and the permutation test on amino acid frequencies were conducted by An et al. (An, et al., 2018). It was showed that Ala (A), Gly (G), Met (M), Arg (R), Val (V), occurred less frequently in type IV effectors than in cytoplasmic proteins. Meanwhile, Phe (F), Ile (I), Lys (K), Asn (N), Ser (S), Tyr (Y), Thr (T) occurred more frequently in type IV effectors than in cytoplasmic proteins. Since different benchmark datasets were used, the final results are debatable and incomplete. However, this is the first time to pay attention to the feature named hydrophobicity_PONP930101.G1, which not only corresponds to the amino acid frequency, but also represents the corresponding hydrophilicity. The SHAP summary plot from TreeExplainer (Lundberg, et al., 2020) succinctly displays the magnitude, prevalence, and direction of a feature’s effect. Each dot in Figure 3(B) corresponds to a protein sample in the study. The position of the dot on the x-axis is the impact that feature has on the model’s prediction for that protein. For example, the higher value of hydrophobicity_PONP930101.G1 has higher contribution on predicting a protein being an effector. In contrast, when the values of top features such as CS_Freq_SS8 and normwaalsvolume.1.residue0 are high, the corresponding Shapley values are negative driving the model prediction towards non-effector class. Besides, there are many long tails mean features with a low global importance which can yet be extremely important for specific samples. From the analysis above, it is necessary and effective to consider many characteristics at the same time.

**Figure 3.**
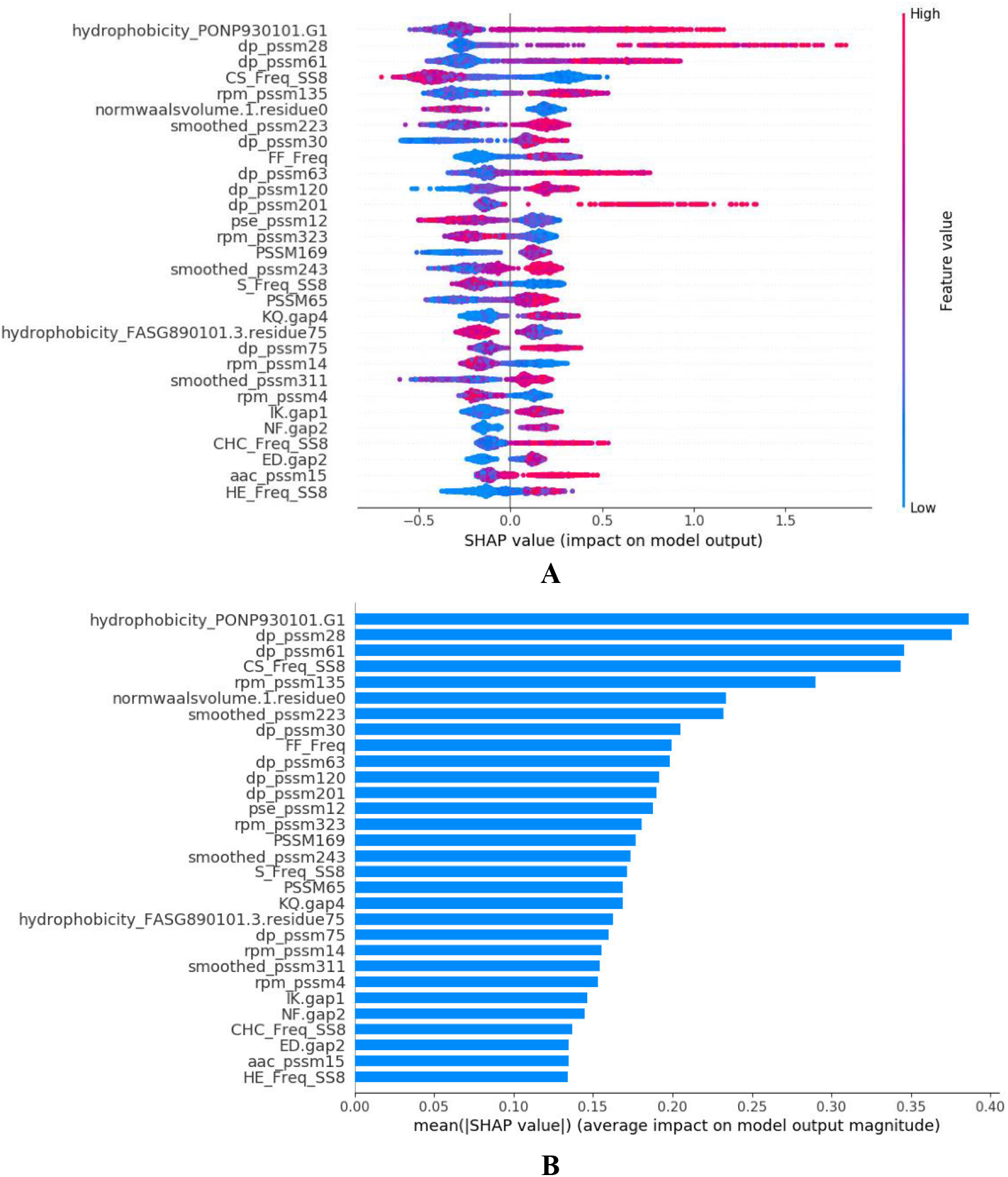
SHAP analysis results for T4SE-XGB. (A) the standard bar plot taken the mean absolute value of the SHAP values for each feature (B) SHAP summary plot sorts features by the sum of SHAP value magnitudes over all samples.

### 4. Conclusion

In this study, we have presented T4SE-XGB, a predictor developed for accurate identification of T4SE proteins based on the XGBoost algorithm. Especially, we have achieved the state-of-the-art performance compared with previous predictors on the benchmark dataset. There are three major conclusions can be drawn. First, compared with different algorithm, the XGBoost algorithm gives more stable and accurate prediction performance for prediction of T4SEs. Second, the feature selection method called ReliefF was utilized to optimize feature vectors, which extracted important features from a large number of candidate features and improved the model performance. Furthermore, unlike other sequence-based T4SEs predictors, T4SE-XGB can provide meaningful explanation based on samples provided using the feature importance and the SHAP method. It gives us the details about how some features, such as DP-PSSM features and hydrophobicity_PONP930101.G1 from CTD contributed to the final direction of prediction. Meanwhile, it explains the reason why it is essential to pay attention to some certain identities, and also considers a variety of features at the same time.

The final results showed that T4SE-XGB achieved satisfying and promising performance which is stable and credible. However, the model is still constrained by the quantity of T4SE proteins which need to be further updated and the characteristics of T4SEs which need to be discovered. Besides, some potential relationships between features need to be explored. In the future, we plan to find and extract as many features as possible from a large amount of collected data to discriminate type IV secreted effectors from non-effectors.

